# Probing the Molecular Interactions of A22 with Prokaryotic Actin MreB and Eukaryotic Actin: A Computational and Experimental Study

**DOI:** 10.1101/2024.04.27.591468

**Authors:** Anuj Kumar, Samiksha Kukal, Anusha Marepalli, Saran Kumar, Sutharsan Govindarajan, Debabrata Pramanik

## Abstract

Actin is a major cytoskeletal system that mediates the intricate organization of macromolecules within cells. The bacterial cytoskeletal protein MreB is a prokaryotic actin-like protein governing cell shape and intracellular organization in many rod-shaped bacteria including pathogens. MreB stands as a target for antibiotic development, and compounds like A22 and its analogue, MP265, are identified as potent inhibitors of MreB. The bacterial actin MreB shares structural homology with eukaryotic actin, despite lacking sequence similarity. It is currently not clear whether small molecules that inhibit MreB can act on the eukaryotic actin due to their structural similarity. In this study, we investigate the molecular interactions between A22 and both MreB and eukaryotic actin through molecular dynamics approach. Employing MD simulations and free energy calculations with an all-atom model, we unveil robust A22-MreB interaction and substantial binding affinity with eukaryotic actin. Experimental assays reveal A22’s toxicity to eukaryotic cells, including yeast and human glioblastoma cells. Microscopy analysis demonstrates profound effects of A22 on actin organization in human glioblastoma cells. Overall, this integrative computational and experimental study advances our understanding of A22’s mode of action and highlights its potential as a versatile tool for probing actin dynamics and as a candidate for therapeutic intervention in pathological conditions like cancer.

## INTRODUCTION

Molecular organization is a fundamental biological process. The intracellular arrangement of macromolecules within the cell is primarily facilitated by dedicated cellular organizing systems such as the cytoskeletal systems. Actin and actin-like cytoskeletal systems are prominent among them^1^. Current evidence suggests that they are universally present across prokaryotes to eukaryotes, occasionally including viruses^2–4^. Actin-based cytoskeletal systems perform a diverse array of functions including cell division, cell migration, intracellular transport, signalling, and regulation of cell shape. Given their pivotal roles, there is significant interest in developing therapeutic drug molecules that specifically target these systems, especially in diseased conditions, with the aim of modulating cellular processes for therapeutic benefit^5,6^. However, a deeper mechanistic understanding of the protein-ligand interactions of actin and actin-binding molecules is pivotal for effectively targeting these systems.

The bacterial actin-like cytoskeletal protein MreB is highly conserved across rod-shaped bacteria^7^. It performs various functions, including cell shape maintenance, intracellular organization, molecular transport, and chromosomal segregation^8–10^. MreB polymerizes in the presence of ATP or GTP, forming double filaments assembled in an antiparallel orientation. In the ATP-bound state, MreB undergoes conformational rearrangement, facilitating nucleotide hydrolysis and depolymerisation^11–13^. Although the role of hydrolysis in polymerization dynamics remains unclear, the dynamic organization of the MreB filaments is well understood to mediate cell wall synthesis and cell shape maintenance in bacterial cells^9,11,14^.

Due to its significance, MreB is considered a crucial target for antibiotic development. A22 (S-(3,4-dichlorobenzyl) isothiourea) and its analog MP265 (4-chlorobenzyl carbamimidothioate) are two well-studied small molecules known to inhibit MreB^15–19^. Various studies have elucidated the interaction of these ligands with the protein and their resultant effects. The presence of these inhibitors converts the cell shape from rod to spherical due to the inhibition of MreB. Initially, A22 was thought to be a competitive inhibitor of the nucleotide ATP (adenosine triphosphate) and could directly bind to the nucleotide-binding pocket of MreB, disrupting the bacterial cytoskeleton^20^. However, subsequent structural studies revealed that A22 can bind near the nucleotide-binding pocket of MreB simultaneously with ATP^11^. Current evidence suggests that A22 affects ATP-Induced conformational change in MreB resulting in destabilization and depolymerisation^21^. Overall, it is evident that A22 affects MreB polymerization leading to filament instability, cell wall defects, and eventual cell death^22^.

MreB shares similarities with eukaryotic actin in terms of filamentous structure formation and is considered a structural homolog of actin^23,24^. However, there are certain differences between the higher-order structures of MreB and actin. For instance, eukaryotic actin filaments consist of two twisted and parallel protofilaments, while bacterial MreB filaments consist of two straight and antiparallel protofilaments^11^. Thus, MreB and actin exhibit significant structural similarity despite lacking significant sequence similarity. Currently, no studies have evaluated whether MreB-inhibiting molecules can also inhibit actin or vice versa.

Actin-mediated processes are highly important during cancer progression, as they promote cell migration and malignant properties of cancer. Some of the hallmarks of cancer cells include altered actin organization, increased expression of actin, and presence of distinct isoforms. For these reasons, actin is considered a drug target for cancer therapeutics, and several inhibitors of actin are developed as potential anticancer agents^25,26^.

To develop a profound knowledge of protein functions and to identify or develop better therapeutic drug molecules for treating various diseases, a thorough mechanistic understanding of protein-ligand interactions is required. By employing computational techniques at the all-atom level, we can explore microscopic information about such biologically relevant molecules. With the parallelization of programming codes and advancements in high-performance computing, we can now perform all-atom level molecular simulations for even macromolecular systems, including explicit water molecules. Using classical all-atom molecular dynamics simulations and integrated biased sampling techniques, various studies have reported a deeper understanding of thermodynamics, kinetics, and dissociation pathways for biological molecules^27–30^, as well as interactions between ligands and proteins, and between nanomaterials and biomolecules^31,32^.

In this study, we explored the ligand-protein interactions of A22 with MreB and eukaryotic actin through a molecular dynamics approach. Employing integrated all-atom molecular dynamics and enhanced sampling approach, we show that A22 can bind with actin with significant binding energy. We further provide experimental validation that A22 affect cell growth in eukaryotes, extending to both yeast and human glioblastoma cells. We provide direct evidence that A22 affects actin organization in glioblastoma cells, highlighting its potential as a promising candidate for cancer therapy.

## METHODS

### COMPUTATIONAL INVESTIGATION

#### System preparation

To model the protein-ligand system, we started with the experimental crystal structure for MreB and Actin proteins. For MreB protein we took Cc-MreB (PDB ID: 4CZG)^33^ and Yeast-Actin (PDB ID: 1YAG)^34^ from the PDB database. We removed the chemical entities which were not required for our studies from the crystal structure using Discovery studio^35^. The missing residues and atoms in the crystal structure were modelled using the swiss-model server^36^ for both proteins. The following portion of the amino acid residues were missing in the MreB protein, and we modelled this specific sequence forming a loop as follows 226ALA, 227ASP, 228GLY, 229GLU and missing atoms in 40HIS, 41GLN, 43ILE, 45VAL, and 111LYS residues in Actin protein. The A22 initial structure (shown in Figure 1) was extracted from Cc-MreB (4CZG) crystal structure. To obtain the optimized structure (as shown in Figure 1), we performed quantum optimization using Gaussian software (Gaussian16)^37^. Initially, we performed geometry optimization at the PM6 level and later M06 level theory, with 6-311G (d,p) basis set and water dielectric medium as continuum model. Subsequently, we calculated single point resp charges^38^ to get the atomic charges of A22 molecule. Starting with the gaussian output as input to the swiss – param server^39^ we developed the Force Field parameters of A22 ligand. We used Open babel software^40^ to the conversion of file format from one into another. To prepare the ligand – protein complex, we place the ligand A22 in the binding pocket of MreB near to the active binding site (NT binding pocket) with a similar conformation as in the crystal structure. Similarly, we prepared the initial complex for the Actin – A22 as well.

**Figure 1:**
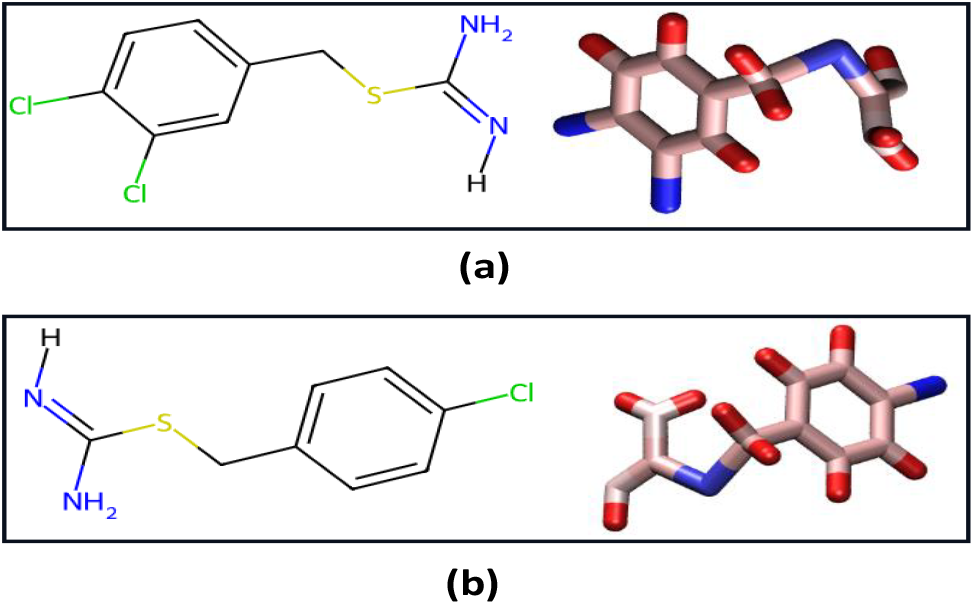
Figure showed 2-D and 3-D structures of the A22 and MP265 ligands. (a) 2-D and 3-D structures of A22, (b) 2-D and 3-D structures of MP265. The 3-D structures are shown in Licorice representation.

#### Molecular dynamics

Starting with the initial protein – ligand complexes, we first solvate the system with water molecules. We used TIP3P^41^ water model for our study. To charge neutralize the system and maintain 0.1 mol/lit. ionic concentration, we add NaCl counterions by substituting water molecules (instantaneous snapshot of initial solvated system is shown in Figure 2). Periodic boundary conditions were applied in all directions to take care of the bulk properties.

**Figure 2:**
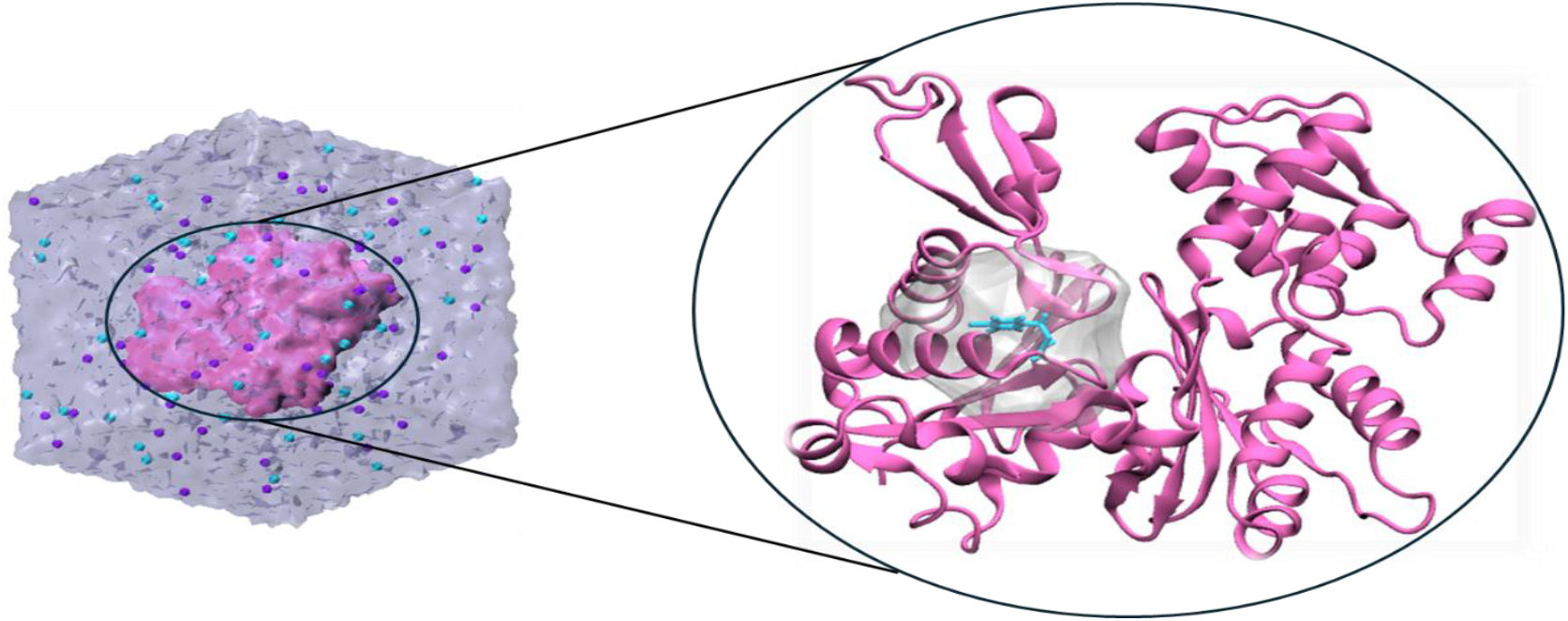
Figure shows the initial prepared system with protein (MreB), ligand (A22), water, and ions (left). The zoomed one shows the initial orientation of ligand in protein binding pocket.

Full system details of the unbiased molecular dynamics simulations have been provided in the supplementary information Table S1. The GROMACS-2022 software package was utilised as MD engine to perform the molecular dynamics simulations at the all-atom level. We modelled the protein interactions using Charm36 force field^42^. To remove the bad contacts among the water molecules with the solute, the energy minimization was carried out by using steepest descent algorithm. The Particle-Mesh Ewald (PME) method was utilized to calculate the long-range electrostatic interaction. Verlet cut-of scheme was used with value of 1.4 nm for short range electrostatic and van der walls interactions. Further Velocity rescale thermostat^43^ has been used to maintain the system temperature at 300 K and Parrinello-Rahman barostat^44^ to maintain the system pressure at 1 atmospheric condition. Once the system is equilibrated, we performed production run. We used leapfrog integrator with an integration time step of 2fs. To constraint the fastest motion associated with the hydrogen atoms in the system, we used LINCS algorithm. The production simulations were performed in isothermal-isobaric (NPT) ensemble. To measure the structural stability and various stable interactions of the systems, we performed long unbiased MD simulations (100ns) and calculated various structural quantities.

To explore the possible three-dimensional interactions between the protein-ligand complexes at the bound state, we calculated the interaction fingerprints. We used online tool, ProLIF, a python library to evaluate the fingerprint interaction. In ProLIF to compute the interaction fingerprint, two groups of atoms (one from protein and another from ligand) were selected using SMART queries. The interaction is determined based on geometrical constraint, distances and /or angles^45^. We computed interaction fingerprints in a binary vector format to understand the nature of protein-ligand intermolecular interactions that remain stable for more than 70% over the 100 ns simulation time. We also showed the protein -ligand interaction network in 2-D form. The way in which protein-ligand interactions are typically represented is residues as nodes, and interactions as edges. All the resultant interactions are the results from the aggregation of all the frame. The frequency with which an interaction is seen control the width of the corresponding edge.

#### Well-tempered Metadynamics

Since dissociation of ligands from the protein binding pocket occurs in longer time scale, to observe dissociation events within reasonable time scale and with our available computational resources, we further performed enhanced sampling simulation for the protein – ligand system. Among various enhanced sampling techniques available, the metadynamics^46^ is a well-established and highly used technique employed for diverse rare event systems, for example various physical^47^, biological^48,49^, chemical^50^, and materials^51^ systems etc. The PLUMED package^52,53^ was integrated with MD engine GROMACS to perform the metadynamics simulation. In a metadynamics simulation, we bias the system on the fly along some pre-defined reaction coordinate, and the system samples the phase space and eventually escape from the minima. By adding the bias/hills we can extract the free energy of unbinding of the ligand from the protein binding pocket. For our study we used well – tempered metadynamics (WT-MetaD)^54^, a variant of metadynamics (MetaD) simulation for all the systems. In a well – tempered metadynamics, the height of the bias is gradually tuned through a bias factor which helps the system to converge faster. We used a gaussian bias with bias height of 2 kJ, width 0.5, bias factor 15, hills deposition frequency 1 ps (every 500 steps), and at temperature 300K. In the WT-MetaD, the system was biased along a pre-defined reaction coordinate. The reaction coordinate is defined as the low dimensional predictor of the system which helps the system to overcome the barrier. It can be defined as the linear or non-linear combination of various order parameters of the system. Here in our study, we used centre of mass distances between two groups (ligand heavy atoms, and protein residues surrounding the binding pocket) as the reaction coordinate (Figure 3a shows the schematic representation of the two group of atoms, and Figure 3b shows two dummy points). For MreB+A22, Actin+A22, MreB+MP265, and Actin+MP265 systems, we performed more than 17 independent simulations for all the systems (23, 19, 21, and 17 independent WT-MetaD runs respectively for MreB+A22, Actin+A22, MreB+MP265, and Actin+MP265) (in supplementary information we provided system details for the well – tempered metadynamics simulations in Table S2).

**Figure 3:**
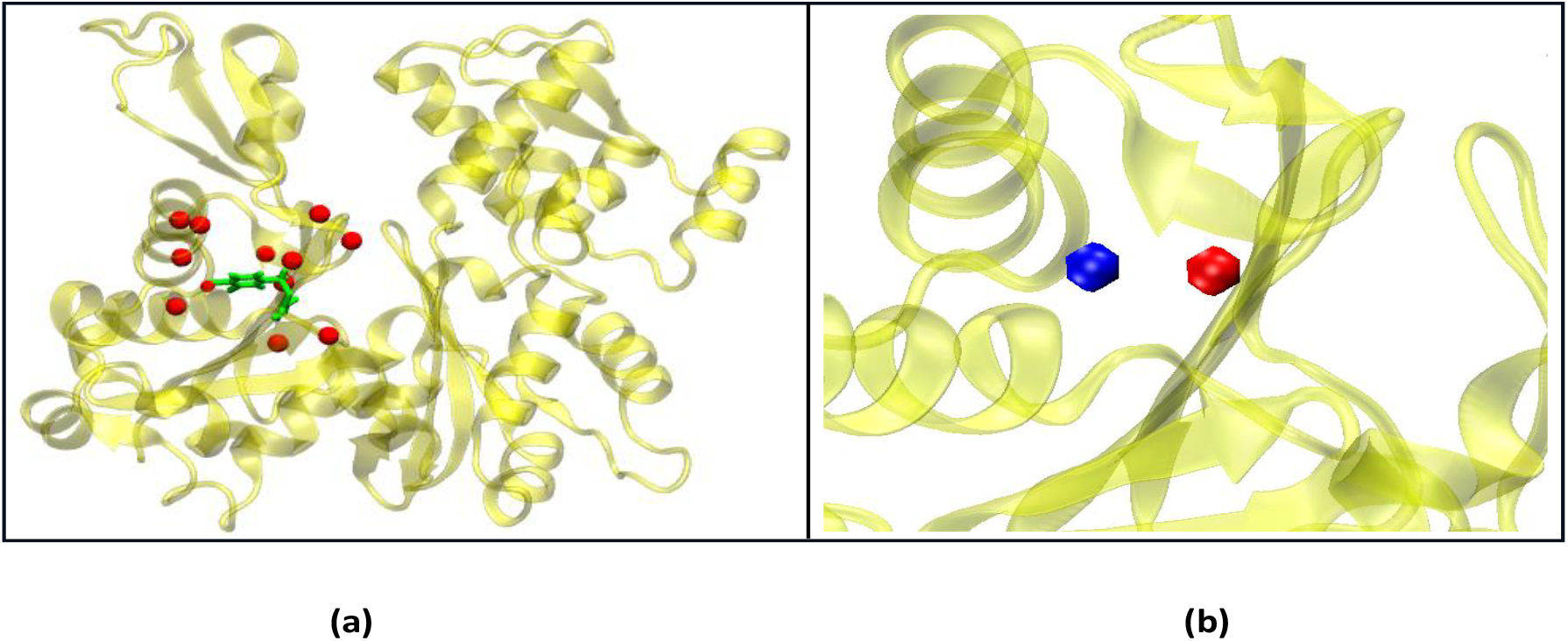
Center of mass between A22 ligand and surrounding protein residues taken as reaction coordinate. A22 (green) and surrounding amino acid residues (red).

To get the free energy we first use plumed plugin sum-hills. Then we do reweighting^55^ to get the free energy profile along the reaction coordinate. From the individual free energy, we calculated probability p = exp(-F/ k_B_T) and then by averaging over probabilities we got averaged free energy F = - k_B_Tln(p). The error bars were calculated from the standard deviation over the independent runs.

### EXPERIMENTAL INVESTIGATION

#### Strains and growth media

*E. coli* MG1655 and *S. cerevisiae* strain BY4741 were used in this study. Overnight *E. coli* cultures were grown in LB at 37°C and overnight *S. cerevisiae* cultures were grown in MM4 medium at 37°C. Antibiotics were added at indicated concentrations. Antibiotics Ampicillin, Kanamycin, and Chloramphenicol were purchased from Sisco Research Laboratories. A22 hydrochloride was purchased from Sun-Shine Chemical (Catalog No: 186211).

#### Spot titer assay

For spot titer assays of *S. cerevisiae* BY4741, overnight cultures grown in MM4 liquid medium were serially diluted 10-fold in fresh MM4 medium. Subsequently, 4 μl of serially diluted samples are spotted onto the surface of the agarized MM4 medium supplemented with A22 or other antibiotics at indicated concentrations. The plates were then incubated at 30°C for 2 days, and images were captured using the Gel Documentation System from Bio-Rad. For spot titer assays of *E. coli* MG1655, LB medium was used instead of MM4 medium and the plates were incubated overnight before imaging.

#### Cell Culture

The cell line used in this work (U87GFP) was purchased from American Type Culture Collection (ATCC, Manassas, VA, USA). U87GFP cell line was grown in DMEM-F12 medium, supplemented with 10% FBS, 1% l-glutamine, 100 U/mL penicillin/streptomycin, and 1% non-essential amino acids (NEAA). U87GFP cell line was maintained at 37°C in a humidified atmosphere of 95% air and 5% CO_2_, and periodically screened for contamination.

#### MTT assay

The cytotoxicity effect of A22 drug on U87GFP cells was evaluated using 3-(4,5-dimethylthiazol-2-yl)-2, 5-diphenyltetrazolium bromide (MTT) assay. Briefly, cells were seeded on 96-well plate for 24 h and subsequently treated with either water or varied concentrations of A22. After 24 h of treatment, cells were washed with 1X PBS and 100 µl MTT (final concentration of 0.5 mg/ml) was added to each well. Incubation with the MTT was done for 3 h at 37 C, following which the formed formazan crystals were dissolved in DMSO. After placing the plate at 37°C for 15 min at dark, absorbance was measured at 570 nm with a reference filter of 630 nm.

#### Phalloidin staining and microscopy

Following drug treatment, U87GFP glioblastoma cells were fixed in 1% paraformaldehyde at room temperature (RT) for 20 min. After washing cells with 1X PBS, cells were permeabilized using 0.2% triton X for 15 min. Non-specific binding was blocked using 3% BSA for 30 min at RT. Cells were stained with Alexa Fluor 647 Phalloidin-iFluor™ 647 Conjugate (Cayman chemical; catalogue: 20555) for 45 min at RT in the dark to stain the F-actin. After washing extra dye once with PBS, the coverslips were mounted with Mobi-oil. Imaging was performed using the Olympus FV1200 confocal microscope.

## RESULTS

### Molecular dynamics simulations and structural stability analysis

We carried out molecular dynamics simulations to delve into the structural stability of protein-ligand interactions. We performed molecular dynamics simulations of ligand A22 with MreB protein (PDB ID: 4CZG) and Yeast-Actin (PDB ID: 1YAG). Starting with equilibrated systems, we conducted 100 ns unbiased all-atom molecular dynamics simulations for all the systems. To verify whether the systems were properly equilibrated, various equilibrium properties were calculated, shown in Figure S1 and Figure S2 for MreB+A22 and Actin+A22 systems, respectively. In Figure 4a, the root mean square deviation (RMSD) of the protein backbone is depicted for MreB/A22, Actin/A22, MreB/MP265, and Actin/MP265 systems. Initially, from 0 ns to 30 ns, there were changes in RMSD for all the systems, but later, RMSD stabilized over the simulation time.

**Figure 4:**
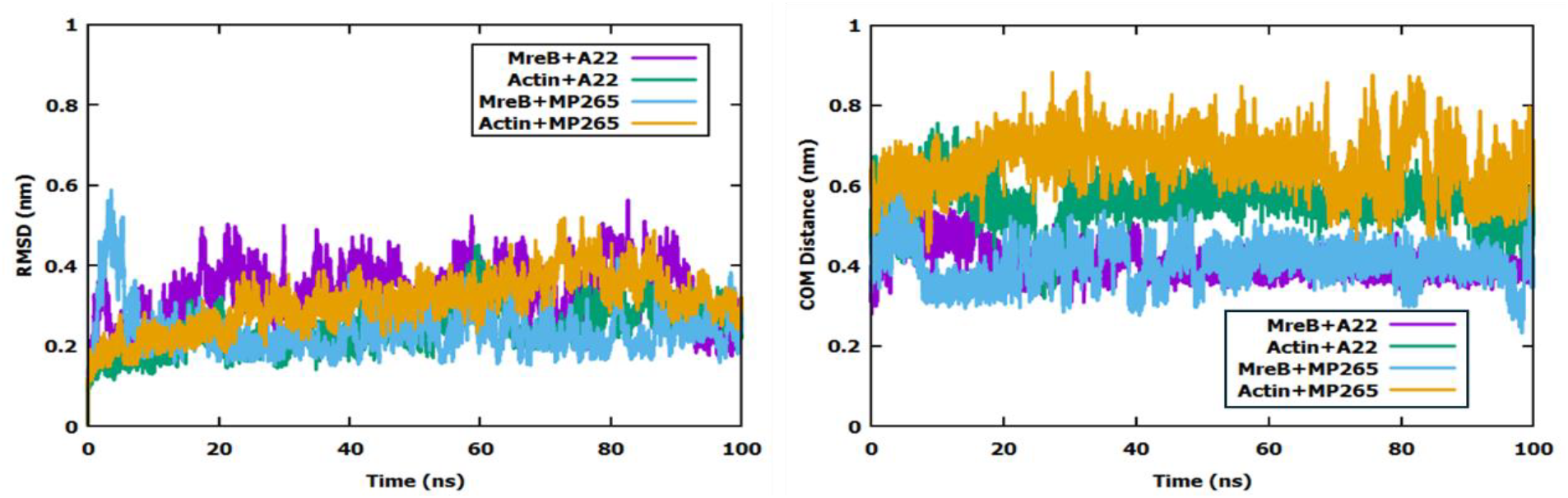
(a) The root mean square deviation (rmsd) of the protein backbone is shown for MreB + A22 (violet), Actin +A22 (green), MreB+MP265 (sky-blue), and Actin+MP265 (yellow) systems, respectively. (b) The COM distance between ligand and protein is shown for MreB + A22 (violet), Actin+A22 (green), MreB+MP265 (sky-blue), and Actin+MP265 (yellow) systems.

To confirm the stability of the ligand in the protein binding pocket, we selected the surrounding pocket residues, identified ligand heavy atoms, and calculated the center of mass for both. We then observed how the center of mass distance varied with time. As depicted in Figure 4b, in the case of the Actin+A22 system, for the first 20 ns, the A22 center of mass distance increased by 0.5 nm, after which it became stable. In contrast, in the MreB+A22 system, the center of mass distance changed by 0.5 nm after 20 ns, stabilized from 20 to 35 ns, then decreased by 0.5 nm again and remained stable for the entire 100 ns simulation. In the MreB+MP265 system, significant fluctuations were observed, but overall, the ligand remained stable. Similarly, in the Actin+MP265 system, the center of mass distance fluctuated by large amounts, demonstrating the dynamic nature of protein-ligand interactions within the pocket. Relatively short deviations in the center of mass distance between the protein and ligand indicate close proximity and stability of the ligand in the pocket, ensuring strong interactions between the protein and ligand.

### Free Energy surface

To explore the binding affinity of the ligand with the protein binding pocket, we observed the dissociation of the ligands from the protein binding pocket and calculated the average dissociation free energy. First, we calculated independent free energies for these four systems and then calculated the average free energies (the independent free energy plots are shown in Figure S3, Figure S4, Figure S5 and Figure S6, respectively). In Figure 5, we depicted the average free energy profiles for all the systems along with their respective error bars. The free energy profile indicates that ligand A22 interacting with protein MreB has a lower free energy (-35.9±2 k_B_T) compared to Actin with A22 (-12±2.5 k_B_T). This indicates a stronger binding of A22 to the MreB protein compared to Actin. From the free energy profile for MP265, we observe that it exhibits lower free energy with MreB (-45.5±2.5 k_B_T) compared to Actin (-15.8±2.5 k_B_T). Hence, we deduce that both A22 and MP265 bind strongly with the MreB protein than Actin. We present the quantitative values of dissociation free energy and the potential interactions in Table 2.

**Figure 5:**
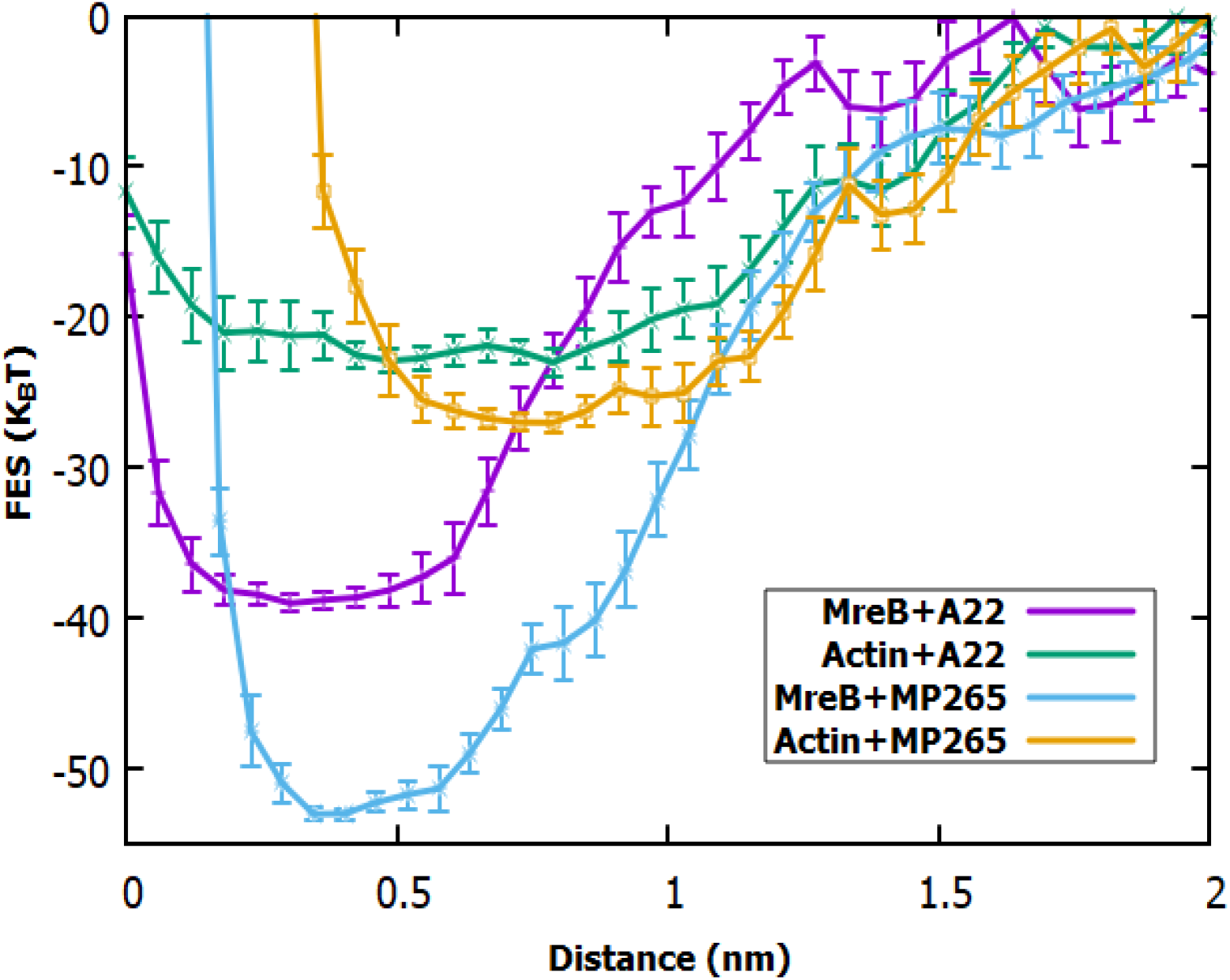
Profiles show of average free energy with distance for MreB + A22 (violet), Actin +A22 (green), MreB+MP265 (sky-blue), Actin+MP265 (yellow), respectively.

### Interactions Fingerprints & 2D Interactions

The examination of interactions fingerprints and two-dimensional interactions provides insights into the robust binding between the A22 ligand and MreB protein. Understanding the origin of such robust binding is crucial. To elucidate the dominant interactions within protein-ligand systems, we conducted an extensive analysis of stable interactions present. Utilizing the 100 ns unbiased MD simulation trajectory, we calculated the interactions fingerprint following the protocol outlined in the methodology section. Figure 6 illustrates all the possible intermolecular interactions observed throughout the entire 100 ns simulation. Across all cases - MreB+A22, Actin+A22, MreB+MP265, and Actin+MP265 - hydrophobic (blue), hydrogen bond (orange & green), and van der Waals (brown) interactions were present over 70% of the time throughout the simulation, as shown in Table 1.^30^Illustrative representations (Figure 6a for MreB+A22 and Figure 6b for Actin+A22) showcase the pivotal residues engaged in ligand interactions. In MreB, residues LEU17, ILE79, PRO112, and ALA115 contribute to hydrophobic interactions, while THR19 and MET74 participate in van der Waals interactions with ligand A22. Additionally, LEU17 forms a hydrogen bond. Conversely, in Actin, VAL76 and ALA108 are involved in hydrophobic interactions, while GLY74, ASN111, and VAL76 engage in van der Waals interactions with the ligand (as shown in Figure 6b).

**Table 1:** It shows the stable interactions (occurrence probability ≥ 70%) of A22 and MP265 with specific residues of MreB and actin.

| Ligand | Protein residues |  | Type of Interactions | Occurrence Probability |
| --- | --- | --- | --- | --- |
| A22 | MreB | LEU17 | Hydrophobic | 0.759824 |
|  |  | LEU17 | HBond | 0.728827 |
|  |  | THR19 | VdW | 0.783822 |
|  |  | MET74 | VdW | 0.926007 |
|  |  | ILE79 | Hydrophobic | 0.987401 |
|  |  | PRO112 | Hydrophobic | 0.992901 |
|  |  | ALA115 | Hydrophobic | 0.873813 |
| A22 | Actin | GLY74 | VdW | 0.805719 |
|  |  | VAL76 | Hydrophobic | 0.974303 |
|  |  | VAL76 | VdW | 0.772423 |
|  |  | ALA108 | Hydrophobic | 0.994001 |
|  |  | ASN111 | VdW | 0.783822 |
| MP265 | MreB | LEU17 | Hydrophobic | 0.932007 |
|  |  | LEU17 | HBond | 0.889911 |
|  |  | LEU17 | VdW | 0.961304 |
|  |  | MET74 | Hydrophobic | 0.772623 |
|  |  | ILE79 | Hydrophobic | 0.978402 |
|  |  | PRO112 | Hydrophobic | 0.993101 |
| MP265 | Actin | ASN12 | Hydrophobic | 0.822618 |
|  |  | ASN12 | VdW | 0.717128 |
|  |  | ILE71 | Hydrophobic | 0.970803 |
|  |  | VAL76 | Hydrophobic | 0.916908 |
|  |  | ALA108 | Hydrophobic | 0.945605 |
|  |  | GLN137 | VdW | 0.714629 |

**Table 2:** It shows the dissociation free energy and number of dominant interactions of specific types for all four systems.

| System | $\Delta G$ (kT) | Number of Interactions |
| --- | --- | --- |

|  |  | H-Bond | VdW | Hydrophobic |
| --- | --- | --- | --- | --- |
| MreB+A22 | $-35.9 \pm 2.0$ | 1 | 2 | 4 |
| Actin+A22 | $-12.1 \pm 2.5$ | 0 | 3 | 2 |
| MreB+MP265 | $-45.5 \pm 2.5$ | 1 | 1 | 4 |
| Actin+MP265 | $-15.8 \pm 2.5$ | 0 | 2 | 4 |

**Figure 6:**
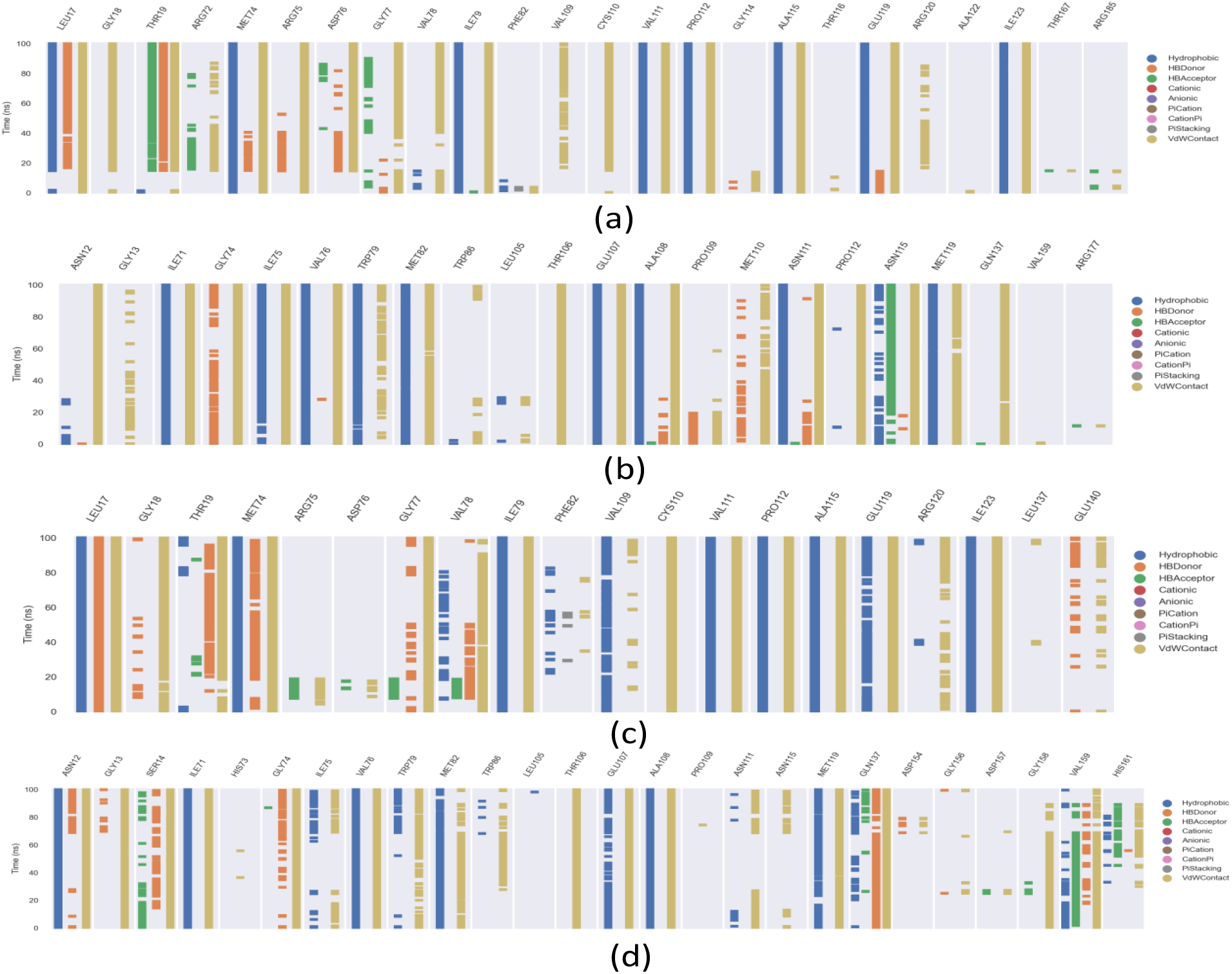
Various kinds of interaction are shown in interaction fingerprints as shown in the Figure (a) MreB + A22, (b) Actin +A22 (c) MreB+MP265, (d) Actin+MP265 respectively.

In Figure 6c, for MreB+MP265, notable hydrophobic interactions involve LEU17, MET74, ILE79, and PRO112 residues, with LEU17 also forming a hydrogen bond. In Actin, MP265 forms hydrophobic interactions with ASN12, ILE71, VAL76, and ALA108, while ASN12 and GLN137 contribute to van der Waals interactions (as shown in Figure 6d). In Figure S7, we show the 2D interaction network plots for (a) MreB+A22, (b) Actin +A22, (c) MreB+MP265 and (d) Actin+MP265 systems, respectively.

Our findings align with previous research conducted by Elvis Awuni et al.^21^, which also identified key residues involved in the interaction between A22 and MreB. These residues include LEU17, THR19, MET74, ILE79, VAL111, PRO112, GLU119, ILE123, and GLU140. Notably, in its apo form, A22 forms a hydrogen bond with LEU17 but not with GLU140, a pattern supported by our results. Specifically, our data indicates that six residues - LEU17, THR19, MET74, ILE79, PRO112, and ALA115 - primarily facilitate the binding of MreB and A22.

#### A22 inhibits growth of yeast and human glioblastoma cells

Our computational analysis suggests that the MreB-inhibiting antibiotic A22 is capable of binding to eukaryotic actin. Since actin is an integral part of eukaryotic cell organization and is important for their survival, we next tested the effect of A22 on eukaryotic cell survival. For this purpose, we first tested the growth of the yeast *S. cerevisiae* by spotting serial dilutions on MM4 agar plates containing A22 at various concentrations and compared it with the growth of *E. coli* in the presence of A22. As expected, A22 affected growth of *E. coli* effectively even at a lower concentration of 5 µg/ml (Figure 7a). In the case of yeast, A22 did not affect cell growth even at 50 µg/ml. However, in the presence of a 100 µg/ml concentration of A22, yeast cells exhibited a 10^5^-log reduction in growth (Figure 7b). In contrast, yeast cells treated with even higher concentrations of other antibiotics, including ampicillin, chloramphenicol, and gentamicin, did not show any inhibitory effect (Figure S8). This suggests that the growth inhibition observed in yeast in the presence of A22 is specific. We also tested whether A22 displays a cytotoxic effect on mammalian cells, using U-87 MG human glioblastoma cell line through MTT assay. Results presented in Figure 7c show that A22 significantly decreased the viability of U-87 MG cells at concentrations above 10 µg/ml, and the IC_50_ value was observed to be around 22 µg/ml. Taken together, the results presented above suggest that A22 exhibits cytotoxic effects on eukaryotic cells.

**Figure 7:**
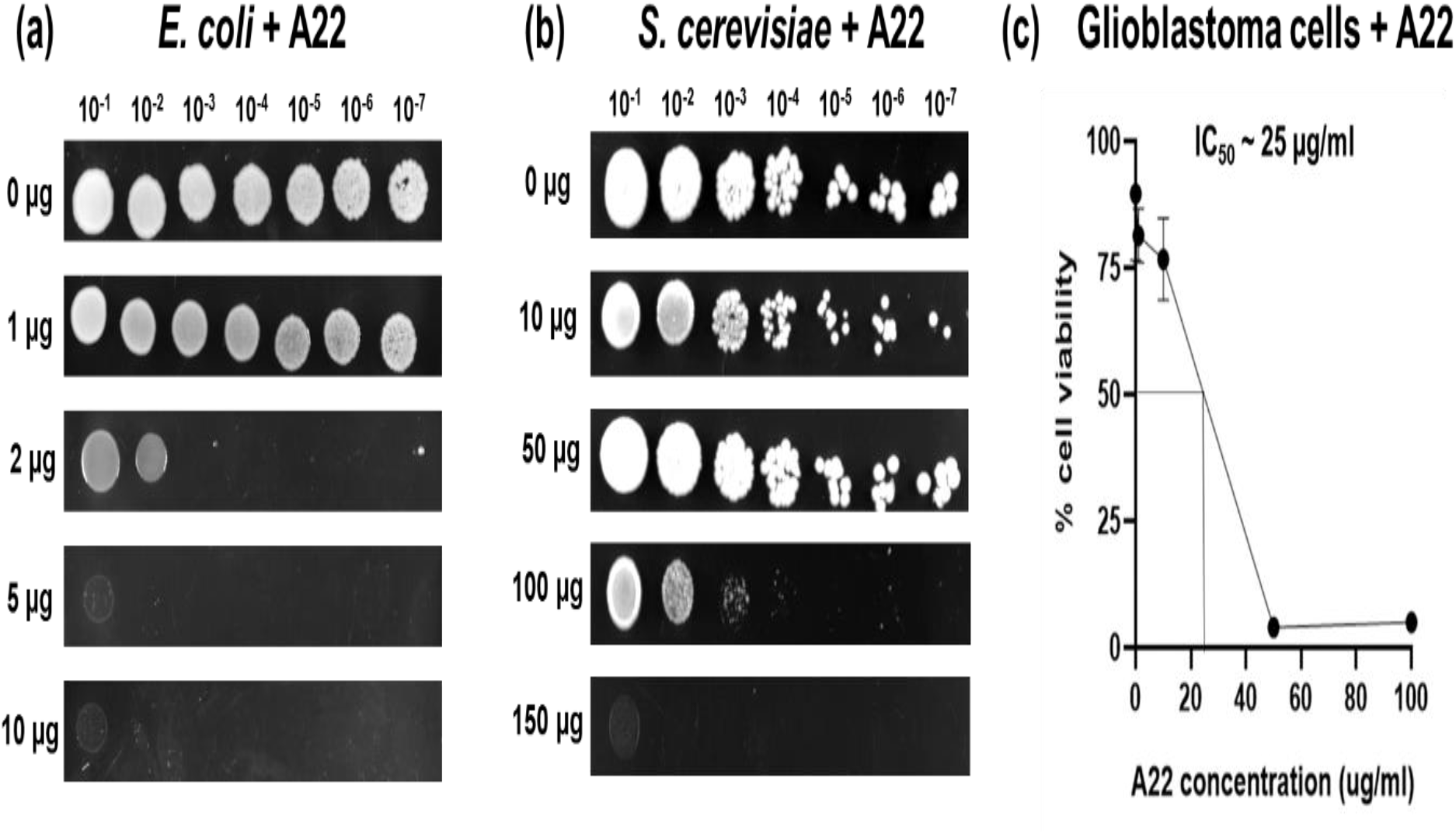
Serial dilutions of *E. coli* (a) or *S. cerevisiae* (b) were performed in the presence of various concentrations of A22. Growth defects were observed as reductions or lack of colony formation (c) Graph shows the percentage of cell viability of the U-87 MG human glioblastoma cell line treated with different concentrations of A22 using the MTT assay. The IC_50_ value is identified as approximately 25 µg/ml.

#### A22 disrupts actin organization

Next, we aimed to elucidate the effect of A22 on actin organization. To achieve this, U-87 MG cells expressing GFP were treated with 20 µg/ml of A22, a concentration close to the IC50 value, and subsequently prepared for actin imaging using phalloidin staining. Phalloidin is well known for its specific binding to and stabilization of filamentous actin (F-actin). Fluorescently labelled phalloidin is a common tool in fluorescence microscopy, facilitating the visualization of actin filaments within cells^56^. Untreated cells exhibited a typical actin localization pattern, characterized by well-defined F-actin formation. Conversely, A22-treated cells displayed a pronounced disorganization of the actin cytoskeleton (Figure 8). We observed contraction of actin filaments, accompanied by a notable decrease in actin fluorescence intensity, which was reduced to 41% compared to the untreated control (Figure S9).

**Figure 8:**
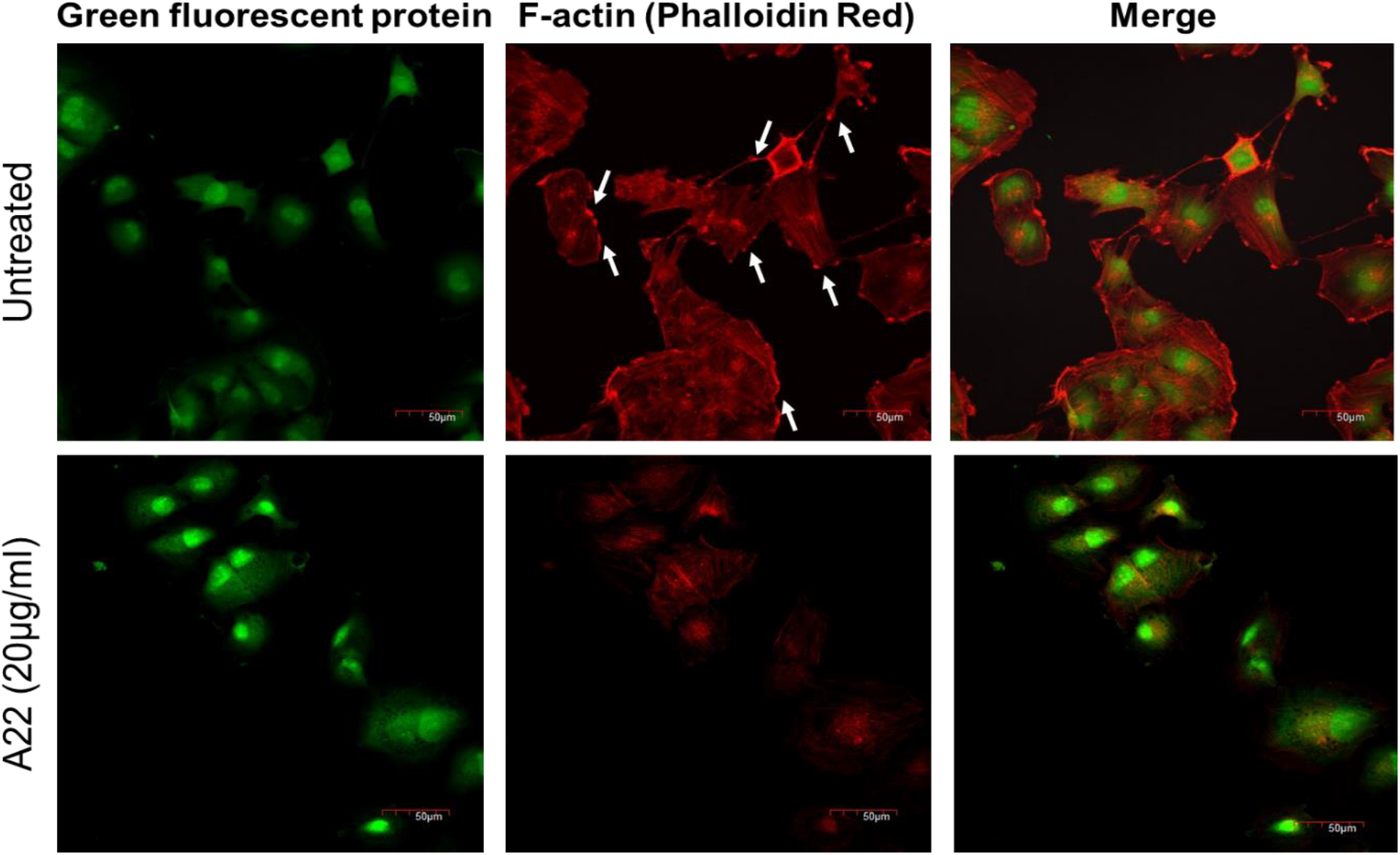
Confocal laser scanning microscopy of phalloidin stained U-87 MG human glioblastoma cell line without treatment (upper panel) or with A22 (20 µg/ml). White arrow indicates F-actin localization. U-87 MG cells express GFP (green), and F-actin was stained with Phalloidin-iFluor™ 647 Conjugate. Scale bar corresponds to 50 µm.

## DISCUSSION

Actin cytoskeletal filaments are fundamental structural component of cells. They play vital roles including regulation of cell division, signalling, motility, and morphological controls. Although they are ubiquitously present from prokaryotes to eukaryotes, they are much more complex in terms of structure, function, and isoforms in eukaryotes^3,9,57,58^. Actin is considered as an important drug target. For example, in bacteria, the MreB actin-like protein is important for virulence, motility, and cell wall stability. Small molecule inhibitors disrupting MreB cytoskeletal protein have potent antibacterial properties against major rod-shaped pathogens including *P. aeruginosa, K. pneumonia, Salmonella* and pathogenic *E. coli* ^17,18,59^. In the case of eukaryotes, actin mediates tumor-relevant processes, including cell migration, invasion, metastasis, and signal transduction pathways implicated in tumorigenesis^17^. Therefore, actin inhibitors are considered as promising candidates as anti-cancer drugs^25^. However, unlike microtubule inhibitors like Ixabepilone and Paclitaxel, which are well established as anti-cancer agents^60^, actin inhibitors are not widely used so far due to poor understanding of the drug-target interactions.

In this study, we explored the structural landscape of MreB – the prokaryotic actin, and the eukaryotic actin and investigated the ability of MreB inhibiting small molecules like A22 to inhibit eukaryotic actin through combined computational and experimental approaches. This investigation is inspired by the fact that MreB and actin share significant structural homology, although their sequences are different^9^. Through MD simulations and free energy calculations with an all-atom model, we show that A22 exhibits strong interaction with MreB. This is not surprising since A22 is well established antibiotic that inhibits MreB^11,61^. Interestingly, we observed significant binding affinity of A22 with eukaryotic actin. Thus, the binding free energy of A22 with MreB is -35.9 ± 2 k_B_T and the binding free energy of A22 with Actin is -12 ± 2.5 k_B_T. Free energy represents the energy difference between the bound state of the (ligand bound to the protein) and the unbound state (ligand and protein separate). In general, a higher negative free energy indicates a stronger binding interaction, suggesting that the formation of the ligand-protein complex is energetically favored. In our case, the more negative binding free energy value for A22 with MreB (-35.9 k_B_T) compared to A22 with Actin (-12 k_B_T) indicates that A22 binds more strongly to MreB than to Actin. Similar trend was also observed for MP265 which exhibited lower binding free energy with MreB (- 45.5±2.5 k_B_T) with respect to actin (-15.8±2.5 k_B_T). Hence, we observe that A22 and MP265, both bind strongly with MreB protein but also significantly with the eukaryotic actin.

Molecular dynamics (MD) simulations are extensively utilized computational methods for identifying and mapping drug-target interactions, as well as for identifying key residues involved in these interactions. This approach has been employed in numerous studies, including some of our previous studies, for uncovering information on drug-target interactions^62–65^. In our investigation, we employed a similar MD simulation approach to elucidate the key amino acid residues of MreB and actin that interact with the drug A22, aiming to understand the nature of these interactions and their potential biological effects. In the case of MreB, we identified five amino acids contributing hydrophobic interactions in the binding interface, with additional contributions from hydrogen bonds (two amino acids) and van der Waals forces (eight amino acids). Importantly, these amino acid sites do not correspond to the ATP binding site of MreB. This is consistent with the observation that A22 is not a competitive inhibitor of ATP, and both molecules can bind simultaneously^11^. Intriguingly, our data did not reveal any known MreB mutations that confer resistance to A22. It is plausible that the amino acids identified in our study as interacting with A22 represent crucial residues for the functioning of MreB. Mutations in these regions might result in lethality, which could explain the absence of isolated resistant mutations. Regarding actin, A22 interacted with only a few amino acids, forming hydrophobic bonds with two amino acids and van der Waals interactions with three amino acids. Although the biological implications of this interaction region are not clear, the lesser interaction of A22 with actin compared to MreB possibly explains the weaker binding affinity towards actin.

Experimental assays revealed that A22 affects cell growth in eukaryotic cells, as assessed in two different and diverged model systems: the yeast S. cerevisiae and mammalian cells, specifically human glioblastoma cells. In human glioblastoma cells, A22 was cytotoxic at concentrations above 10 µg/ml, with an IC50 value of nearly 22 µg/ml. Intriguingly, previous experiments have shown that A22 treatment does not induce genotoxic effects against human peripheral blood mononuclear cells^17^. The reason for the discrepancy is not clear, but the same study observed A22-induced growth defects at higher concentrations. Thus, it is possible that permeability and exposure time could explain the observed differences. In yeast cells, A22 exhibited toxicity at higher concentrations ranging up to 100 µg/ml. However, unlike A22 exposure, when yeast cells were treated with other bacterial antibiotics like ampicillin, chloramphenicol, or gentamycin, no growth defects were observed, even after exposure to very high concentrations. This, along with other observations, suggests that these eukaryotic cells are specifically affected by A22, indicating the presence of A22 binding targets, potentially actin. Indeed, microscopy analysis revealed profound effects of A22 on actin organization in human glioblastoma cells. This was evidenced through phalloidin staining, which is a widely used technique in cell biology to visualize and study F-actin structures within cells^56^. The observed delocalization of actin by A22 is reminiscent of other well-established actin inhibitors like Cytochalasin D^66^. Cytochalasins interact with actin filaments by capping the barbed ends, thereby directly altering filament assembly and dynamics^67^. However, the nature of A22’s interaction with actin and its effects on actin dynamics, filament assembly, and disassembly dynamics are yet to be explored. Biochemical and biophysical assays using purified actin and A22, and high-resolution fluorescence microscopy observation of actin dynamics in the presence of A22 will provide detailed information on the nature of A22-actin interactions. These aspects will be investigated in our future studies.

Actin contributes to cancer progression by increasing cell motility, invasiveness, and metastatic potential. Cancer cells also exhibit other actin-related hallmarks, including aberrant actin isoform expression, actin-mediated chemoresistance, and altered signal transduction^26,68^. Due to these aspects, actin inhibitors are explored as anti-cancer agents. Our experiments with human glioblastoma cells indicate that A22 exhibits anti-cancer activity; however, whether this property is specific to cancer cells needs further study. Our future studies will investigate the therapeutic potential of A22 and the possibility of using A22 as a valuable research tool for investigating actin biology.

In conclusion, our integrated computational and experimental approach provides comprehensive insights into the structural stability, binding affinity, and functional implications of A22-mediated targeting of the major prokaryotic actin MreB and eukaryotic actin. These findings contribute to the understanding of cytoskeletal organization and offer avenues for the development of multipurpose therapeutic agents with potential clinical applications.

## Supplementary Information

System details for unbiased molecular dynamics and well-tempered metadynamics simulations (Table S1 and S2); equilibrium properties of the MreB–A22 and Actin– A22 complex (Figure S1 and S2), independent free energy profiles for MreB – A22, Actin – A22, MreB – MP265, Actin – MP265 (Figure S3, S4, S5, S6), respectively; 2D interaction plots for these four systems (Figure S7); serial dilutions of *S. cerevisiae* in presence of antibiotics - Ampicillin (Amp), Chloramphenicol (Cm), and Gentamycin (Gm) (Figure S8); mean fluorescent intensity of phalloidin stained U-87 MG human glioblastoma cells in untreated vs A22-treated group (Figure S9) have been provided in the supplementary information.

## Acknowledgments

A.K and A.M thanks SRM University – AP for PhD fellowship. We thank the HPCC computing facilities provided by SRM University – AP supercomputer centre. DP thanks Science and Engineering Research Board (SERB), Government of India for supporting with the State University Research Excellence (SURE) project No SUR/2022/004576. S.G acknowledges support from DST-SERB (CRG/2020/003295). Samiksha was supported by IITD PDF fellowship. S.K was funded by grants from DST, DBT and ICMR (SRG/2022/000657, BT/PR46424/MED/30/2449/2023 and IIRP-2023-2341 respectively). We acknowledge Writoban Basu Ball, SRM University-AP, for gift of *S. cerevisiae* BY4741. All authors appreciate the infrastructure support received from SRM University – AP, India (SRMAP/URG/GENERAL/2023-24/010).

## Views

The views expressed in this study are those of the authors and not necessarily those of either the funding agency or any other institution.

## Author contributions

S.G, and D.P formulated the study design and plans. A.K performed the computational experiment. S.K, A.M, S.K performed other experiments. A.K, S.G, and D.P wrote the manuscript.

## References

1. Svitkina, T. M. Actin Cell Cortex: Structure and Molecular Organization. Trends in Cell Biology vol. 30 Preprint at 10.1016/j.tcb.2020.03.005 (2020).

2. Jockusch, B. M. & Graumann, P. L. The long journey: Actin on the road to pro-and eukaryotic cells. Rev Physiol Biochem Pharmacol 161, (2012).

3. Van Den Ent, F., Amos, L. A. & Lo, J. Prokaryotic Origin of the Actin Cytoskeleton. NATURE vol. 413 http://www.nature.com (2001).

4. Da Cunha, V. et al./person-group>. Giant Viruses Encode Actin-Related Proteins. Mol Biol Evol 39, (2022).

5. Rao, J. & Li, N. Microfilament Actin Remodeling as a Potential Target for Cancer Drug Development. Curr Cancer Drug Targets 4, (2005).

6. Giganti, A. & Friederich, E. The actin cytoskeleton as a therapeutic target: state of the art and future directions. Progress in cell cycle research vol. 5 Preprint at (2003).

7. Jones, L. J. F., Carballido-López, R. & Errington, J. Control of cell shape in bacteria: Helical, actin-like filaments in Bacillus subtilis. Cell 104, (2001).

8. Carballido-López, R. The Bacterial Actin-Like Cytoskeleton. Microbiology and Molecular Biology Reviews 70, 888–909 (2006).

9. Shaevitz, J. W. & Gitai, Z. The structure and function of bacterial actin homologs. Cold Spring Harbor perspectives in biology vol. 2 Preprint at 10.1101/cshperspect.a000364 (2010).

10. Govindarajan, S. & Amster-Choder, O. The bacterial Sec system is required for the organization and function of the MreB cytoskeleton. PLoS Genet 13, (2017).

11. van den Ent, F., Izoré, T., Bharat, T. A. M., Johnson, C. M. & Löwe, J. Bacterial actin MreB forms antiparallel double filaments. Elife 2014, (2014).

12. Pande, V., Mitra, N., Bagde, S. R., Srinivasan, R. & Gayathri, P. Filament organization of the bacterial actin MreB is dependent on the nucleotide state. Journal of Cell Biology 221, (2022).

13. Mao, W. et al./person-group>. On the role of nucleotides and lipids in the polymerization of the actin homolog MreB from a Gram-positive bacterium. Elife 12, (2023).

14. Carballido-López, R. & Errington, J. A dynamic bacterial cytoskeleton. Trends in Cell Biology vol. 13 Preprint at 10.1016/j.tcb.2003.09.005 (2003).

15. Iwai, N. et al./person-group>. Structure-activity relationship of S-benzylisothiourea derivatives to induce spherical cells in Escherichia coli. Biosci Biotechnol Biochem 68, (2004).

16. Iwai, N., Nagai, K. & Wachi, M. Novel S-Benzylisothiourea Compound That Induces Spherical Cells in Escherichia coli Probably by Acting on a Rod-shape-determining Protein(s) Other Than Penicillin-binding Protein 2. Biosci Biotechnol Biochem 66, (2002).

17. Bonez, P. C. et al./person-group>. Antibacterial, cyto and genotoxic activities of A22 compound ((S-3, 4 -dichlorobenzyl) isothiourea hydrochloride). Microb Pathog 99, (2016).

18. Awuni, E. Status of Targeting MreB for the Development of Antibiotics. Frontiers in Chemistry vol. 7 Preprint at 10.3389/fchem.2019.00884 (2020).

19. Gitai, Z., Dye, N. A., Reisenauer, A., Wachi, M. & Shapiro, L. MreB actinmediated segregation of a specific region of a bacterial chromosome. Cell 120, (2005).

20. Bean, G. J. et al./person-group>. A22 disrupts the bacterial actin cytoskeleton by directly binding and inducing a low-affinity state in MreB. Biochemistry 48, 4852–4857 (2009).

21. Awuni, E. & Mu, Y. Effect of A22 on the conformation of bacterial actin MreB. Int J Mol Sci 20, (2019).

22. Young, K. D. Why spherical Escherichia coli dies: The inside story. Journal of Bacteriology vol. 190 Preprint at 10.1128/JB.01975-07 (2008).

23. Van Den Ent, F., Amos, L. & Löwe, J. Bacterial ancestry of actin and tubulin. Current Opinion in Microbiology vol. 4 Preprint at 10.1016/S1369-5274(01)00262-4 (2001).

24. Young, G. National Fitness and the New Order. Aust Q 14, (1942).

25. Stehn, J. R. et al./person-group>. A novel class of anticancer compounds targets the actin cytoskeleton in tumor cells. Cancer Res 73, (2013).

26. Suresh, R. & Diaz, R. J. The remodelling of actin composition as a hallmark of cancer. Translational Oncology vol. 14 Preprint at 10.1016/j.tranon.2021.101051 (2021).

27. Payne, R. T., Crivelli, S. & Watanabe, M. All-Atom Simulations Uncover Structural and Dynamical Properties of STING Proteins in the Membrane System. J Chem Inf Model 62, (2022).

28. Galano-Frutos, J. J., Nerín-Fonz, F. & Sancho, J. Calculation of Protein Folding Thermodynamics Using Molecular Dynamics Simulations. J Chem Inf Model 63, (2023).

29. Pramanik, D., Pawar, A. B., Roy, S. & Singh, J. K. Mechanistic insights of key host proteins and potential repurposed inhibitors regulating SARS-CoV-2 pathway. J Comput Chem 43, (2022).

30. Namsani, S., Pramanik, D., Khan, M. A., Roy, S. & Singh, J. K. Metadynamics-based enhanced sampling protocol for virtual screening: case study for 3CLpro protein for SARS-CoV-2. J Biomol Struct Dyn 40, (2022).

31. Pramanik, D. & Maiti, P. K. Dendrimer assisted dispersion of carbon nanotubes: A molecular dynamics study. Soft Matter 12, (2016).

32. Pramanik, D. & Maiti, P. K. DNA-Assisted Dispersion of Carbon Nanotubes and Comparison with Other Dispersing Agents. ACS Appl Mater Interfaces 9, (2017).

33. van den Ent, F., Izoré, T., Bharat, T. A. M., Johnson, C. M. & Löwe, J. Bacterial actin MreB forms antiparallel double filaments. Elife 2014, (2014).

34. Vorobiev, S. et al./person-group>. The structure of nonvertebrate actin: Implications for the ATP hydrolytic mechanism. Proc Natl Acad Sci U S A 100, (2003).

35. Biovia, D. S. Discovery Studio Visualizer v21.1.0.20298. BIOVIA, Dassault Systèmes Preprint at (2005).

36. Arnold, K., Bordoli, L., Kopp, J. & Schwede, T. The SWISS-MODEL workspace: A web-based environment for protein structure homology modelling. Bioinformatics 22, (2006).

37. Frisch, M. J., Trucks, G. W. & H. B. Schlegel. Gaussian 16, Revision C.01. Gaussian. Inc.: Wallingford CT Preprint at (2016).

38. Bayly, C. I., Cieplak, P., Cornell, W. D. & Kollman, P. A. A well-behaved electrostatic potential based method using charge restraints for deriving atomic charges: The RESP model. Journal of Physical Chemistry 97, (1993).

39. Zoete, V., Cuendet, M. A., Grosdidier, A. & Michielin, O. SwissParam: A fast force field generation tool for small organic molecules. J Comput Chem 32, (2011).

40. O’Boyle, N. M. et al./person-group>. Open Babel: An Open chemical toolbox. J Cheminform 3, (2011).

41. Jorgensen, W. L., Chandrasekhar, J., Madura, J. D., Impey, R. W. & Klein, M. L. Comparison of simple potential functions for simulating liquid water. J Chem Phys 79, (1983).

42. Best, R. B. et al./person-group>. Optimization of the additive CHARMM all-atom protein force field targeting improved sampling of the backbone φ, ψ and sidechain χ1 and χ2 Dihedral Angles. J Chem Theory Comput 8, (2012).

43. Bussi, G., Donadio, D. & Parrinello, M. Canonical sampling through velocity rescaling. Journal of Chemical Physics 126, (2007).

44. Parrinello, M. & Rahman, A. Polymorphic transitions in single crystals: A new molecular dynamics method. J Appl Phys 52, (1981).

45. Bouysset, C. & Fiorucci, S. ProLIF: a library to encode molecular interactions as fingerprints. J Cheminform 13, (2021).

46. Laio, A. & Parrinello, M. Escaping free-energy minima. Proc Natl Acad Sci U S A 99, (2002).

47. Martoňák, R. et al./person-group>. Simulation of structural phase transitions by metadynamics. Zeitschrift fur Kristallographie 220, (2005).

48. Pramanik, D., Smith, Z., Kells, A. & Tiwary, P. Can One Trust Kinetic and Thermodynamic Observables from Biased Metadynamics Simulations?: Detailed Quantitative Benchmarks on Millimolar Drug Fragment Dissociation. Journal of Physical Chemistry B 123, (2019).

49. Kaufhold, W. T., Pfeifer, W., Castro, C. E. & Di Michele, L. Probing the Mechanical Properties of DNA Nanostructures with Metadynamics. ACS Nano 16, (2022).

50. Piccini, G., Mendels, D. & Parrinello, M. Metadynamics with Discriminants: A Tool for Understanding Chemistry. J Chem Theory Comput 14, (2018).

51. Martoňák, R., Donadio, D., Oganov, A. R. & Parrinello, M. Crystal structure transformations in SiO2 from classical and ab initio metadynamics. Nat Mater 5, (2006).

52. Bonomi, M. et al./person-group>. PLUMED: A portable plugin for free-energy calculations with molecular dynamics. Comput Phys Commun 180, (2009).

53. Tribello, G. A., Bonomi, M., Branduardi, D., Camilloni, C. & Bussi, G. PLUMED 2: New feathers for an old bird. Comput Phys Commun 185, (2014).

54. Barducci, A., Bussi, G. & Parrinello, M. Well-tempered metadynamics: A smoothly converging and tunable free-energy method. Phys Rev Lett 100, (2008).

55. Tiwary, P. & Parrinello, M. A time-independent free energy estimator for metadynamics. Journal of Physical Chemistry B 119, (2015).

56. Wulf, E., Deboben, A., Bautz, F. A., Faulstich, H. & Wieland, T. Fluorescent phallotoxin, a tool for the visualization of cellular actin. Proc Natl Acad Sci U S A 76, (1979).

57. Löwe, J. & Amos, L. A. Evolution of cytomotive filaments: The cytoskeleton from prokaryotes to eukaryotes. International Journal of Biochemistry and Cell Biology vol. 41 Preprint at 10.1016/j.biocel.2008.08.010 (2009).

58. Esue, O., Cordero, M., Wirtz, D. & Tseng, Y. The assembly of MreB, a prokaryotic homolog of actin. Journal of Biological Chemistry 280, 2628–2635 (2005).

59. Kotzialampou, A., Protonotariou, E., Skoura, L. & Sivropoulou, A. Synergistic Antibacterial and Antibiofilm Activity of the MreB Inhibitor A22 Hydrochloride in Combination with Conventional Antibiotics against Pseudomonas aeruginosa and Escherichia coli Clinical Isolates. Int J Microbiol 2021, (2021).

60. Loong, H. H. & Yeo, W. Microtubule-targeting agents in oncology and therapeutic potential in hepatocellular carcinoma. OncoTargets and Therapy vol. 7 Preprint at 10.2147/OTT.S46019 (2014).

61. Awuni, Y., Jiang, S., Robinson, R. C. & Mu, Y. Exploring the A22-Bacterial Actin MreB Interaction through Molecular Dynamics Simulations. Journal of Physical Chemistry B 120, 9867–9874 (2016).

62. Wang, Y., Lupala, C. S., Liu, H. & Lin, X. Identification of Drug Binding Sites and Action Mechanisms with Molecular Dynamics Simulations. Curr Top Med Chem 18, (2018).

63. Smith, Z., Pramanik, D., Tsai, S. T. & Tiwary, P. Multi-dimensional spectral gap optimization of order parameters (SGOOP) through conditional probability factorization. Journal of Chemical Physics 149, (2018).

64. Pramanik, D., Smith, Z., Kells, A. & Tiwary, P. Can One Trust Kinetic and Thermodynamic Observables from Biased Metadynamics Simulations?: Detailed Quantitative Benchmarks on Millimolar Drug Fragment Dissociation. Journal of Physical Chemistry B 123, (2019).

65. Kumawat, A., Namsani, S., Pramanik, D., Roy, S. & Singh, J. K. Integrated docking and enhanced sampling-based selection of repurposing drugs for SARS-CoV-2 by targeting host dependent factors. J Biomol Struct Dyn 40, (2022).

66. Kim, M., Song, K., Jin, E. J. & Sonn, J. Staurosporine and cytochalasin d induce chondrogenesis by regulation of actin dynamics in different way. Exp Mol Med 44, (2012).

67. Cooper, J. A. The role of actin polymerization in cell motility. Annu Rev Physiol 53, (1991).

68. Gibieža, P. & Petrikaitė, V. The regulation of actin dynamics during cell division and malignancy. Am J Cancer Res 11, (2021).

